# Identification of regulatory genes through global gene expression analysis of a *Helicobacter pylori* co-culture system

**DOI:** 10.1101/523274

**Authors:** Nuria Tubau-Juni, Josep Bassaganya-Riera, Andrew Leber, Victoria Zoccoli-Rodriguez, Barbara Kronsteiner, Monica Viladomiu, Vida Abedi, Raquel Hontecillas

## Abstract

*Helicobacter pylori* is a gram-negative bacterium that establishes life-long infections by inducing immunoregulatory responses. We have developed a novel *ex vivo H. pylori* co-culture system to identify new regulatory genes based on expression kinetics overlapping with that of genes with known regulatory functions. Using this novel experimental platform, in combination with global transcriptomic analysis, we have identified five lead candidates, validated them using mouse models of *H. pylori* infection and *in vitro* co-cultures under pro-inflammatory conditions. Plexin domain containing 2 *(Plxdc2)* was selected as the top lead immunoregulatory target. Gene silencing and ligand-induced activation studies confirmed its predicted regulatory function. Our integrated bioinformatics analyses and experimental validation platform has enabled the discovery of new immunoregulatory genes. This pipeline can be used for the identification of genes with therapeutic applications for treating infectious, inflammatory, and autoimmune diseases.

## Introduction

Chronic bacterial infections trigger complex and dynamic host-bacterial interactions that modulate immunometabolic host responses. The phenotypic manifestation of these dynamic interactions results from the coordinated expression of blocks of genes with overlapping functions that cooperate in modulating host responses. In addition to pathogen-associated molecular patterns (PAMPs) that signal infection and are associated with induction of innate anti-bacterial and inflammatory responses, other lesser known bacterial components elicit compensatory immunoregulatory responses that when exploited by pathogens, promote bacterial persistence. For instance, *Mycobacterium tuberculosis* induces IL-10-driven regulatory responses that suppress the activated immune mechanisms and contribute to long-term infection (Gong et al., 1996; Moreira-Teixeira et al., 2017; O’Leary, O’Sullivan, & Keane, 2011; Redford, Murray, & O’Garra, 2011). Therefore, the activation of host immunoregulatory mechanisms by certain bacterial organisms inhibit the effector immune response and prevent bacterial clearance.

*Helicobacter pylori* is a gram-negative, microaerophilic, spiral-shaped bacterium with unipolar, sheathed flagella (Kusters, van Vliet, & Kuipers, 2006; O’Rourke & Bode, 2001) that constitutes the primary member of the gastric mucosa in infected individuals (Noto & Peek, 2017; Sheh & Fox, 2013). *H. pylori* is highly specialized to colonize the human gastric niche. The infection is chronic and affects more than 50% of the world’s population (Hooi et al., 2017). *H. pylori* infection is mostly asymptomatic; however, approximately 10% of carriers will develop peptic ulcers (Ernst & Gold, 2000; Wroblewski, Peek, & Wilson, 2010), and 1-3% gastric cancer (Wroblewski et al., 2010). Interestingly, *H. pylori* infection may be an important driver of systemic tolerance in asymptomatic individuals with an inverse correlation between the presence of this bacterium and the development of autoimmune diseases, asthma, esophageal adenocarcinoma and type-2-diabetes (Bassaganya-Riera et al., 2012; de Martel et al., 2005; Reibman et al., 2008; van Wijck et al., 2018; Xie et al., 2013). These conflicting implications may stem from the relative predominance of antagonistic immune responses that encompass both effector and regulatory components elicited by *H. pylori (Bhuiyan et al., 2014; Kabisch, Semper, Wustner, Gerhard, & Mejias-Luque, 2016; Raghavan & Quiding-Jarbrink, 2012; Smythies et al., 2000).* However, long-term colonization by the bacterium, due to the failure of the immune system to clear the infection, suggests that the strong *H. pylori-induced* regulatory responses can shift inflammatory/effector responses leading to chronicity of the infection.

Macrophages have been described as key immune cells in *H. pylori-induced* regulatory mechanisms (Leber et al., 2016). Particularly, *H. pylori* interacts with a specific subset of mononuclear phagocytes that generate IL-10-driven regulatory responses facilitating optimal colonization of the gastric mucosa (Viladomiu et al., 2017). We demonstrated that macrophage peroxisome proliferator-activated receptor gamma (PPARγ), an anti-inflammatory transcription factor, was needed for the induction of the full spectrum regulatory response (Viladomiu et al., 2017). Additional macrophage-expressed genes *(Par1, HO-1)* have been shown to play a similar role to PPARγ and contribute to keeping high levels of colonization while reducing pathology and disease (Chionh et al., 2015; Gobert et al., 2014).

Early after the initiation of the immune response, both macrophages and dendritic cells (DC) endure strong metabolic and transcriptional changes. To increase the speed of energy production and facilitate the execution of effector responses, activated macrophages are subjected to a metabolic switch, in which glycolysis and lactate production predominate, while Krebs cycle and oxidative phosphorylation are reduced into secondary roles (Kelly & O’Neill, 2015; Rodriguez-Prados et al., 2010). This substantial change implies that the metabolic shift towards higher glucose consumption and glycolysis rate is key to enable the required spectrum of host responses in inflammatory macrophages. In contrast, alternatively-activated macrophages present a metabolic component in which oxidative phosphorylation is dominant and fatty acid consumption is increased (Jha et al., 2015; Kelly & O’Neill, 2015; Vats et al., 2006). Moreover, IL-10 was found to suppress glycolysis while stimulating oxidative metabolism (Ip, Hoshi, Shouval, Snapper, & Medzhitov, 2017), suggesting that some metabolic changes are essential for the induction of regulatory responses in macrophages. Interestingly, PPARγ-driven regulatory responses encompass a profound metabolic component since PPARγ is required for glucose and fatty acid uptake, achieve greater fatty acid β-oxidation in macrophages and maintain mitochondrial biogenesis, both dominant processes in alternatively activated macrophages (Odegaard et al., 2007). These metabolic changes are tightly coupled to the suppression of proinflammatory gene expression (Namgaladze et al., 2014; Vats et al., 2006).

Transcriptional changes play crucial roles in modulating immunomodulatory and metabolic responses to infection. Gene expression is highly coordinated in time, with sets of genes with overlapping roles sharing the same expression pattern by being upregulated and downregulated simultaneously. The loss of a single gene can affect the equilibrium of the whole system and have a significant impact in the outcome of the response. Indeed, suppression or inactivation of a single regulatory protein, for instance PPARγ, results in stronger inflammation, while activation or enhanced expression leads to a balanced response, maintained by induction of immunoregulation (Hontecillas et al., 2011 ? Odegaard et al., 2007; Viladomiu et al., 2017; Viladomiu et al., 2012).

In this study, we used an *ex vivo H. pylori* co-culture system to identify genes with putative regulatory function based on the kinetic pattern of expression of known genes using WT and PPARγ-deficient bone marrow-derived macrophages (BMDM). Using a global transcriptomic assay together with a bioinformatics pipeline based on expression pattern-analysis approaches, we have identified five potential new regulatory genes. Extensive *in vitro* and *in vivo* validation studies, in both pro-inflammatory as well as regulatory-induced conditions, support the regulatory functions of the selected group of candidate genes. In particular, the plexin domain containing 2 (Plxdc2) gene was selected as the lead immunoregulatory target, based on its characteristics and expression pattern, for further validation of its regulatory behavior. In conclusion, this manuscript establishes a novel integrated platform for the identification of genes such as *Plxdc2* with promising regulatory and immunometabolic functions that could become new molecular targets to treat inflammatory and autoimmune diseases.

## Results

### *H. pylori* induces expression of regulatory genes in WT but not in PPARγ-deficient macrophages

The *in vivo* interplay between *H. pylori* and the myeloid cell compartment revealed that macrophages are highly responsive to *H.* pylori-induced regulatory responses (Viladomiu et al., 2017). Here, we sought to explore *H. pylori* interactions with macrophages employing a synchronized gentamycin protection co-culture system comparing cells obtained from WT and PPARγ fl/fl;LysCre+ (LysCre+) mice. Gentamycin was applied to the culture system 15 min after cells were exposed to live *H. pylori* to avoid constant extracellular stimulation and synchronize the cellular response. Of note, *H. pylori* can be internalized and replicate in the intracellular compartment (Chu, Wang, Wu, & Lei, 2010; Oh, Karam, & Gordon, 2005; Tang et al., 2012; Wang, Wu, & Lei, 2009). Here, we used intracellular replication post-gentamycin treatment as a marker of the effect and status of the anti-bacterial response. LysCre+ mice lack PPARγ transcription factor in myeloid cells, which results in defective expression of genes with regulatory function and overexpression of pro-inflammatory and antibacterial response genes (Guirado et al., 2018; Malur et al., 2009; Szanto et al., 2010). Cells were harvested at several time-points from 0 to 12 hours post-gentamycin exposure to assess the bacterial burden and changes in gene expression in response to *H. pylori.* The same pattern of bacterial replication was observed for both genotypes. Initial replication was first detected 30 min post-gentamycin treatment and the peak occurred at 120 min (**Figure 1A**). However, bacterial counts in co-cultures of LysCre+ macrophages were significantly reduced throughout the time course, starting at 60 min post-challenge and up to 240 min post-challenge. This phenotype was compatible with an inflammatory shift in LysCre+ BMDM due to the loss of PPARγ resulting from altered activation of certain regulatory *H. pylori-induced* mechanisms and consequently a more efficient anti-bacterial response. To validate this assessment, WT and LysCre+ macrophages were classically activated through LPS/IFNγ stimulation 24 hours prior to the infection. Generation of pro-inflammatory WT macrophages resulted in drastic suppression of *H. pylori* loads at the 120 min time point, compared to the WT control (**Figure 1B**). Moreover, the bacterial burden peak was entirely abrogated in LPS/IFNγ-treated LysCre+ cells. Regarding the *H. pylori-induced* gene expression profile, IFNγ expression following *H. pylori* challenge was remarkably increased in LysCre+ cells compared to WT. The WT group showed a minimal increase at all timepoints relative to time 0 (**Figure 1F**). In contrast, WT macrophages displayed a dramatic increase of the anti-inflammatory cytokine IL-10 at 60 min post-H. *pylori* co-culture, that was significantly diminished in LysCre+ BMDM (**Figure 1E**). Therefore, *H. pylori* promotes the activation of cytokine-driven regulatory mechanisms in WT macrophages, that modulate the immune response and generate a regulatory microenvironment that facilitates bacterial proliferation reaching the highest peak at 2 hours post gentamycin treatment.

**Figure 1.**
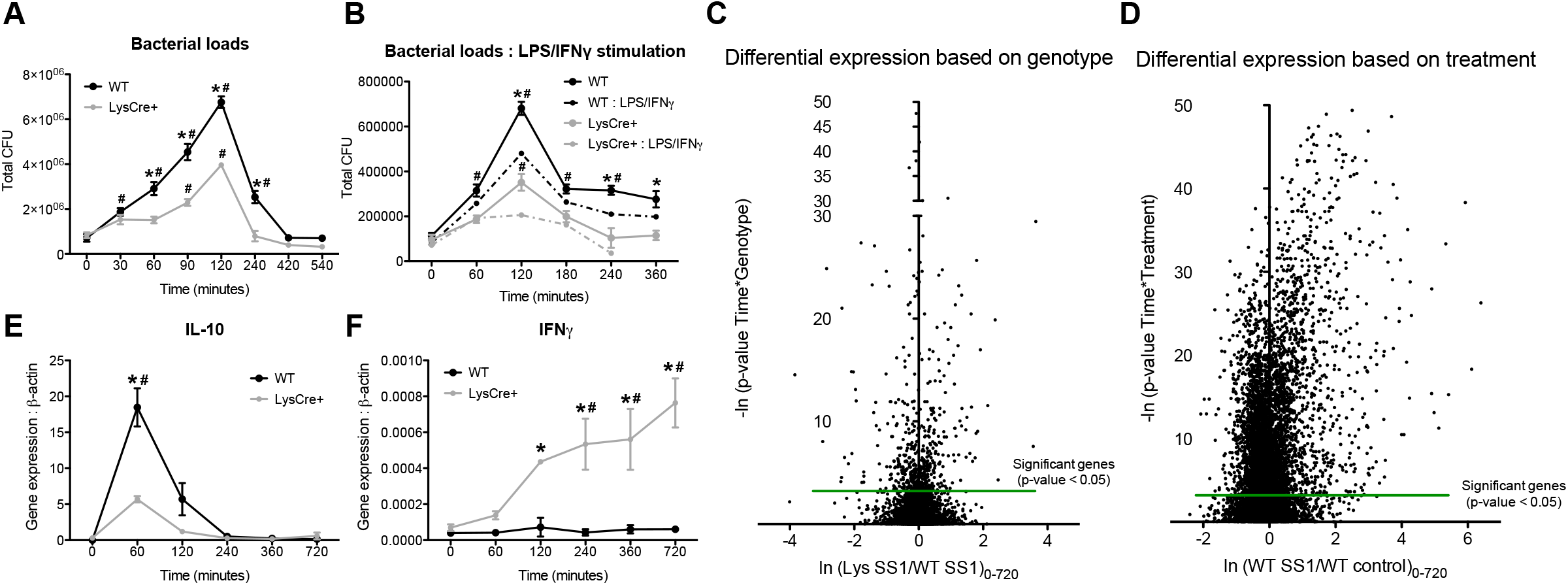
*Helicobacter pylori (H. pylori)* co-culture strongly alters macrophage transcriptomic profile, leading to the activation of early regulatory responses and increasing bacterial persistence in WT cells. WT and PPARγ-deficient (LysCre+) BMDM were co-cultured *ex vivo* with *H. pylori* and cells were harvested at several time-points ranging from 0 to 720 minutes after gentamycin treatment. Bacterial burden **(A)** and gene expression, including IL-10 **(E)** and IFNγ **(F)** were assessed. Differential gene expression based on genotype (C) and *H. pylori* infection **(D)** from a whole transcriptomic analysis performed on the harvested cells were also assessed. To classically activate macrophages prior to *H. pylori* co-culture, cells were stimulated with LPS and IFNγ, then challenged with *H. pylori* and the bacterial burden was measured **(B)**. \**P*-value<0.05 between genotypes, #*P*-value<0.05 within each genotype compared to time 0.

To elucidate the underlying regulatory molecular mechanisms activated upon *H. pylori* co-culture in WT macrophages, we performed a global transcriptomic analysis on a gentamycin protection assay time course (0, 60, 120, 240, 360 and 720 min). RNAseq analysis demonstrated important differences in the gene expression profile within both genotype (**Figure 1C**) and treatment (**Figure 1D**). Almost 50% of genes exhibited a significant differential expression based on the treatment, with a substantial upregulation after *H. pylori* challenge. Thus, *H. pylori* strongly influences the macrophage transcription profile, resulting in drastic modifications in macrophage function that favor the generation of a regulatory phenotype. Therefore, we sought to utilize the new co-culture system to explore novel regulatory pathways activated upon *H. pylori* infection to discover new host regulatory genes that modulate the immune response, with the potential to become molecular targets for the development of therapeutics for infectious, inflammatory and autoimmune diseases.

### Validation of experimental co-culture system with identification of differential expression patterns in characterized antimicrobial genes

Expression of regulatory and pro-inflammatory genes is tightly regulated and coordinated over time. Indeed, these sets of genes frequently present opposite expression kinetics under the same conditions. We selected nine established canonical pro-inflammatory genes to initially explore and identify distinctive patterns of expression associated with pro-inflammatory and antimicrobial functions. Inflammation-related genes are characterized by an increased expression level in *H. pylori-* infected groups compared to the controls (**Figure 2 – Figure supplement 1**), and in the majority of genes, *H.* pylori-induced upregulation is achieved at the later time points of the co-culture (**Figure 2 – Figure supplement 1B-F, H-I**). As expected, lack of PPARγ results in higher gene expression of pro-inflammatory genes. We then performed a 3-way ANOVA analysis that reported the differential expression of eight genes, including *Chil1, Etv5, Iigp1, Ptger4, Sqle, Osm, Rptoros* and *Hspa2* (**Figure 2 – Figure supplement 2**). Most of the genes revealed by the 3-way ANOVA are associated with known pro-inflammatory functions. We focused our analysis in *Chil1* (**Figure 2 – Figure supplement 2A**), *Iigp1* (**Figure 2 – Figure supplement 2C**) and *Sqle* (**Figure 2 – Figure supplement 2E**) due to their expression kinetics that resemble the identified inflammatory genes pattern and their well-known associated role to the host response against pathogens. RNAseq validation through qRT-PCR revealed a similar expression pattern in *Chil1* and *Iigp1,* characterized by a later upregulation due to the infection in both genotypes in case of *ligp1* (**Figure 2E**) or only in infected LysCre+ cells in case of *Chil1* (**Figure 2D**). In contrast, the later induction of *Sqle* following *H. pylori* challenge, was undetectable in the qRT-PCR analysis (**Figure 2F**), which resulted in no differences within groups among the entire time course. In addition, *Chil1* and *ligp1* gene silencing (**Figure 2 – supplement 3A-B**) led to a definite increase of bacterial loads in both genotypes (**Figure 2G**). Indeed, the significant decrease of bacterial burden in LysCre+ macrophages, when compared to the WT group, was abrogated due to *Chil1* or *ligp1* gene knockdown. As expected, there were no differences in culture systems due to silencing of *Sqle.* Therefore, the initial analysis of this global transcriptomics dataset based on patterns of expression related to pro-inflammatory functions highlighted two genes, *Chil1* and *ligp1,* with a relevant, previously established role in macrophage antimicrobial responses. Thus, it validated the potential use of this co-culture system for the identification of novel host immunoregulatory genes by means of kinetic pattern analysis within genotypes and among the time course.

**Figure 2.**
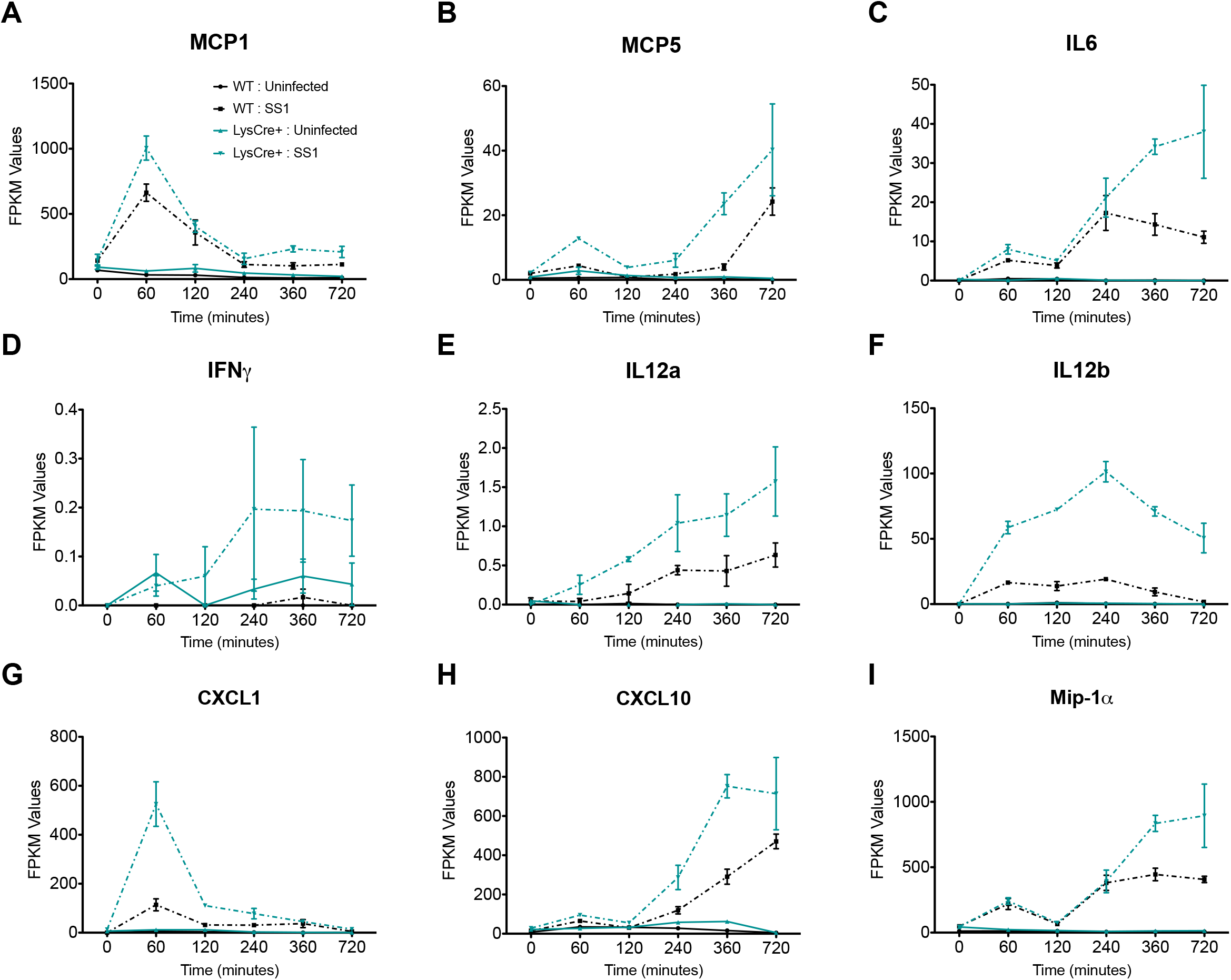
Initial analysis and validation of the whole transcriptomic analysis revealed two genes with differential expression pattern and well-defined anti-microbial functions. Plots represent RNAseq reads of *Chil1* **(A)**, *ligp1* **(B)**, and *Sqle* **(C)** during the entire time-course comparing genotypes and treatments. *Chil1* **(D)**, *ligp1* **(E)**, and *Sqle* **(F)** gene expression from the same co-culture was validated through qRT-PCR. Bacterial loads **(G)** were measured 120 minutes post-gentamycin treatment of *H. pylori* co-cultures in WT and LysCre+ macrophages transfected with *Chil1*-targeted, *ligp1*-targeted or Sqle-targeted siRNA or a scrambled sequence as a negative control. \**P*-value<0.05 between genotypes, #*P*-value<0.05 within each genotype compared to time 0.

### Bioinformatics pattern-expression analysis identified five candidates as novel genes with putative regulatory functions

Nod-like receptors (NLRs) are a subfamily of pattern recognition receptors that like PPARs, regulate innate immune responses and metabolism. Upon ligand binding, downstream activation of the NLR pathway results in the initiation and regulation of potent inflammatory mechanisms, including inflammasome formation and NF-κB activity (Kim, Shin, & Nahm, 2016; Zhong, Kinio, & Saleh, 2013). NLR and PPAR canonical immune pathways were selected to identify the opening set of expression patterns of interests. NLR and PPAR pathways encompass more than 200 genes and modulate the immune response through the activation of transcription and/or induction of metabolic changes in immune cells. Further, even the established role of the NLR pathway in innate immunity activation and the PPAR pathway association to regulatory mechanisms, both canonical pathways inconjunction contain a mixture of genes presenting a dominant pro-inflammatory or anti-inflammatory role, leading to combined expression patterns in the dataset that allows the identification of regulatory patterns.

Initially, we performed an analysis based on the fold change of gene expression between genotypes for each gene of both NLR and PPAR pathways across the entire time course presented in the form of heat maps (**Figure 3A, D**) where blue represents genes downregulated in WT compared to LysCre+, while red represents upregulation of gene expression in WT related to LysCre+. We anticipated the presence of inverted patterns of regulatory and pro-inflammatory genes between both genotypes, where regulatory genes would be increased in WT compared to PPARγ-deficient, and pro-inflammatory genes overexpressed in PPARγ-deficient group. Indeed, the bioinformatics analysis revealed specific expression patterns in both signaling pathways that were clustered in groups. The NLR pathway includes two well-defined clusters based on the distinct genotype gene expression (**Figure 3A**). The orange box, at the top, contains genes upregulated in LysCre+ macrophages, and the green box, at the bottom, contains a second class of genes with greater expression in the WT group. Interestingly, the LysCre+ upregulated genes have a delayed expression pattern, while, WT upregulated genes presented an earlier peak. The two NLR clusters are represented in the PPAR pathway, also depicted in orange and green boxes (**Figure 3D**). The analysis of genes associated with PPAR revealed an additional third cluster, highlighted in purple, including a group of genes characterized by a dysregulated pattern. Particularly, those genes exhibited oscillating expression kinetics in each genotype among the entire time course. A plausible explanation is the existence of a strong PPARγ interaction with these genes. Therefore, the absence of the transcription factor in LysCre+ macrophages could alter the expression of the genes, due to direct activation, inhibition or even due to the upregulation of compensatory mechanisms, that result into a fluctuating expression pattern.

**Figure 3.**
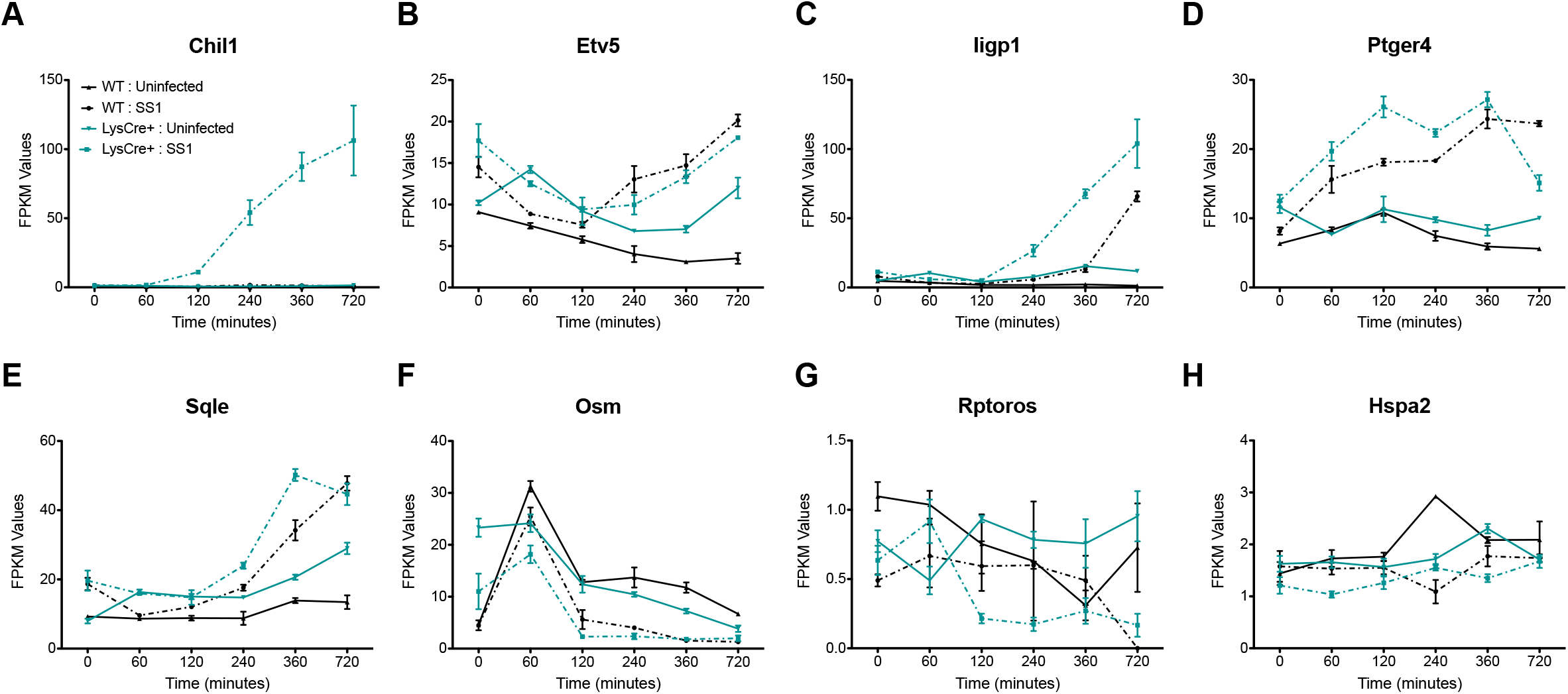
Bioinformatics pipeline utilized to analyze the RNAseq dataset and establish the differential expression patterns that lead to the identification of the potential regulatory candidates. Heatmaps represent the genotype fold-change expression from each gene in the NLR **(A)** and PPAR **(D)** pathways. Blue represents inhibited expression in WT macrophages compared to the upregulation in LysCre+, while red indicates upregulation in WT compared to a suppressed expression in LysCre+ macrophages. NLR **(B)** and PPAR **(E)** pathways are represented in this diagram. In red are highlighted the top genes, based on the differential expression analysis represented in the heatmap, and selected as seed genes. Schematic representation of the steps performed during the bioinformatics analysis, after seed genes selection **(C)**. The 21 genes included in the final set were classified in five groups based on their expression pattern **(F)**.

Gene selection was narrowed to the green box cluster (i.e. genes upregulated in WT), to include genes with a positive response to *H. pylori* in WT macrophages that resembled the peak of bacterial loads reported in this genotype (**Figure 1A**). The choice included 7 NLR (**Figure 3B**) and 10 PPAR (**Figure 3E**) pathway genes highlighted in red, which were defined as seed genes. Based on the expression patterns of the seed genes, we built an initial dataset that comprised both these original genes and a group described as linked genes obtained from the global transcriptome dataset. Linked genes are characterized by their similar expression pattern and linked functions to the seed genes. Certain differentially expressed patterns were selected in this large initial dataset, and through 2 cycles of clustering within the entire global transcriptional dataset, we obtained a specific group of candidates exhibiting the defined expression kinetics. To further narrow down our search, we utilized the Pubmatrix (Becker et al., 2003) tool to select the most novel genes based on the following criteria: number of publications, cell location, and known function (**Figure 3C**). The 21 genes included in the final set were divided in 5 groups based on their expression pattern (**Figure 3 – Figure supplement 1**). Groups 1 and 2 exhibit a clear distinctive pattern within genotype, whereas no genotype differences were observed in groups 3, 4 and 5 (**Figure 3F**). Further, the first two groups exhibit a drastic upregulation post-H. *pylori* challenge in WT that was abrogated in PPARγ-deficient macrophages. Therefore, genes included in groups 1 and 2 displayed an expression pattern potentially associated with a regulatory function.

Based on the pattern analysis, five genes from groups 1 and 2 of the final dataset were identified as potential new regulatory leads for further validation. Plexin domain containing 2 *(Plxdc2,* **Figure 4A**), V-set and immunoglobulin domain containing 8 *(Vsig8,* **Figure 4C**), Ankyrin repeat domain 29 *(Ankrd29,* **Figure 4D**) and C1q and tumor necrosis factor related protein 1 *(Ciqtnf1,* **Figure 4E**) share an early expression peak in WT macrophages, abrogated in LysCre+, that coincides with the bacterial burden spike in the gentamycin protection assay. The kinetics of Protein phosphatase 1 regulatory subunit 3E *(Ppp1r3e,* **Figure 4B**) is slightly different since it was still upregulated in the last timepoint. However, in LysCre+ macrophages, the expression pattern of all five candidates was consistently downregulated and displayed as a flat line. Known properties of these five genes are described in **Table 1**. Publications linked to each of the genes reveal a large diversity of established functions; however, association with the immune system or the immune response was not reported for the majority of the genes. Interestingly, cellular location is also highly heterogeneous. Only two candidates, *Plxdc2* and *Vsig8*, both plasma membrane receptors, have identified ligands. Thus, the limited number of publications together with the current established function of each gene support the novelty of their potential interaction with host immunoregulatory mechanisms.

**Figure 4.**
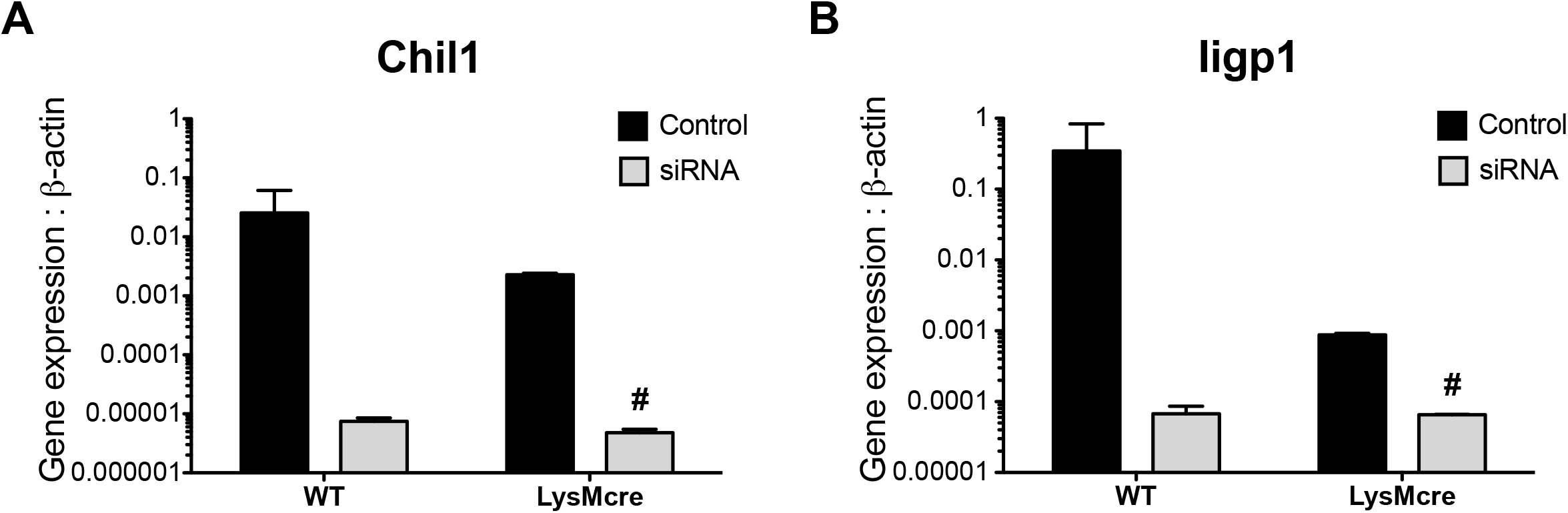
Expression kinetics of the five candidates selected from the bioinformatics analysis to undergo experimental validation. *Plxdc2* **(A)**, *Ppp1r3e* **(B)**, *Vsig8* **(C)**, *Ankrd29* **(D)**, and *C1qtnf1* **(E)** RNAseq reads in WT and LysCre+ macrophages expressed as FPKM values.

**Table 1.** Gene information and properties of the five top selected candidates.

| Gene Symbol | Gene Full Name | Number of publications | Cellular Location | Function/Role | Ligands |
| --- | --- | --- | --- | --- | --- |
| Plxdc2 | Plexin domain containing 2 | 3 (22) | Plasma membrane | PEDF receptor (IL-10 expression).<br>Nervous system development | PEDF |
| Ppp1r3e | Protein phosphatase 1, regulatory (inhibitor) subunit 3E | 1 | Cytoplasm | Glycogen metabolism (increases glycogen synthesis). | None |
| Vsig8 | V-set and immunoglobulin domain containing 8 | 2 | Plasma membrane | Expressed in hair shaft, follicle, nail unit, and oral cavity. VISTA receptor (potential target for cancer or infectious disease) | VISTA (not commercialized) |
| Ankrd29 | Ankyrin repeat domain 29 | 1 | Nucleus | Niemann-Pick Type C diseases (lipid transport) | None |
| C1qtnf1 | C1q and tumor necrosis factor related protein 1 | 24 (27) | Cytoplasm, nucleus | Adipokyne related to glucose metabolism. Link to Pparγ (increased expression with rosiglitazone treatment). | None |

### *In vivo* and *in vitro H. pylori-induced* upregulation of all five lead target candidates in WT mice is abrogated in pro-inflammatory macrophages from PPARγ null mice

To perform a validation of the five selected genes, we initially measured their expression, by qRT-PCR on samples from each time point of the gentamycin protection assay used in the global transcriptomic analysis. Similar to the observed pattern in the RNAseq dataset, *H. pylori* co-culture induces a significant early upregulation of all five candidates in WT macrophages starting at 60 minutes post co-culture. After 60 minutes, the expression of *Plxdc2* (**Figure 5A**), *Vsig8* (**Figure 5E**), *Ankrd29* (**Figure 5G**) and *C1qtnf1* (**Figure 5I**) begin to decline in cells obtained from WT mice to reach same levels as the LysCre+ BMDM at 360 or 720 minutes. Similar to the results from the RNAseq data, *Ppp1r3e* (**Figure 5C**) expression was maintained at high levels throughout the time course analysis. Cultures from LysCre+ BMDM exposed to *H. pylori* failed to upregulate expression of the selected genes. Therefore, there is an early upregulation of the five genes in WT BMDM that was suppressed in cells with a pro-inflammatory phenotype due to the loss of PPARγ.

**Figure 5.**
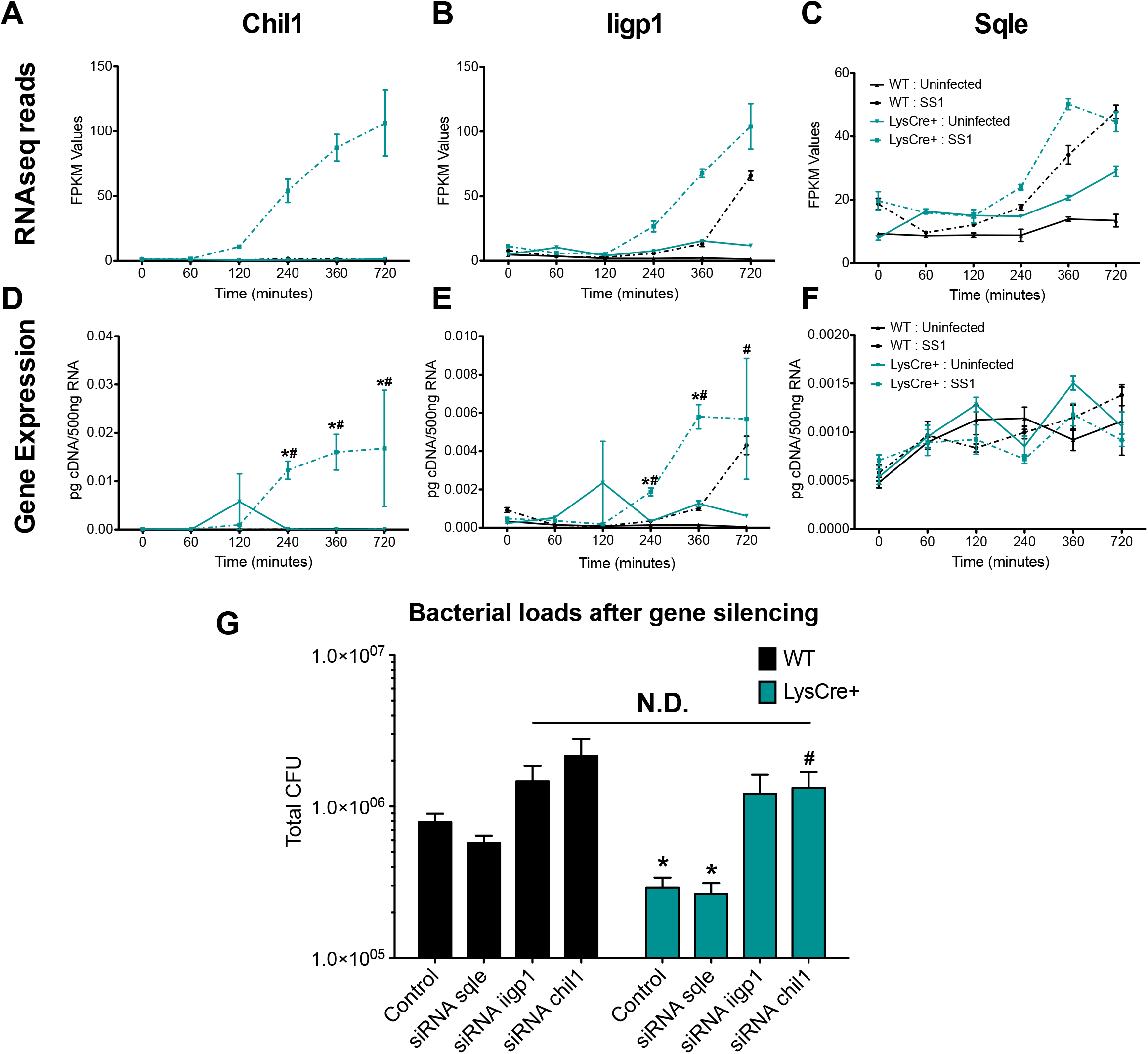
*In vivo* and *in vitro* validation of the five selected candidates under *Helicobacter pylori-induced* regulatory conditions. WT and LysCre+ BMDM were co-cultured *ex vivo* with *H. pylori* and harvested at several time-points ranging from 0 to 720 minutes. *Plxdc2* **(A)**, *Ppp1r3e* **(C)**, *Vsig8* **(E)**, *Ankrd29* **(G)**, and *C1qtnf1* **(I)** gene expression was measured by qRT-PCR. WT and LysCre+ mice were infected with *H. pylori*SS1 strain. Non-infected mice were used as control. *Plxdc2* (B), *Ppp1r3e* **(D)**, *Vsig8* **(F)**, *Ankrd29* **(H)** and *C1qtnf1* (J) gene expression was measured by qRT-PCR. \**P*-value<0.05 between genotypes, #*P*-value<0.05 within each genotype compared to time 0. For in vivo experiments, n=4.

To explore the dynamics of the selected genes *in vivo,* WT and LysCre+ mice were infected with *H. pylori.* In a previous study, we demonstrated that *H. pylori* infection *in vivo* induces strong regulatory mechanisms driven by IL-10-expressing myeloid cells starting at day 14 post-infection and reaching maximum levels at day 28 post-infection (Viladomiu et al., 2017). To evaluate whether induction of regulatory responses *in vivo* alters the kinetics of targeted genes, stomachs were collected at day 28 from infected and non-infected mice. Consistent with the *in vitro* findings, *H. pylori* infection upregulated the expression of *Plxdc2* (**Figure 5B**), *Ppp1r3e* (**Figure 5D**), *Vsig8* (**Figure 5F**), *Ankrd29* (**Figure 5H**) and *C1qtnf1* (**Figure 5J**) in WT gastric tissue. In contrast, minimal or no differences were reported upon *H. pylori* infection in PPARγ-deficient mice. Therefore, activation of regulatory responses after *H. pylori* challenge in WT mice correlate with increased transcription of the five selected genes. However, *H. pylori* infection under pro-inflammatory conditions, due to the lack of PPARγ, abrogates this effect on the selected lead regulatory genes.

### Activation of pro-inflammatory responses modulate the dynamics of the top lead regulatory target genes

To further characterize the potential regulatory functions of the selected genes, we sought to assess their behavior under inflammatory conditions in a controlled environment *in vitro.* Briefly, WT and LysCre+ BMDM were treated with 100ng/mL of LPS for 60, 120, 240, 360 and 720 minutes. LPS administration *in vitro* activates BMDM and modulates their cytokine profile. Particularly, LPS upregulates TNFα expression with a significant increment in LysCre+ macrophages (**Figure 6F**). In contrast, the IL-10-induced peak reported in WT macrophages is abrogated by the lack of PPARγ (**Figure 6H**). Additionally, LPS treatment suppressed PPARγ expression starting at 60 minutes post challenge (**Figure 6G**). The results show a slight decrease in *Plxdc2* (**Figure 6A**), *Vsig8* (**Figure 6C**) and *C1qtnf1* (**Figure 6E**) expression after LPS treatment plus a significant downregulation of *Ppp1r3e* starting at 120-minutes post stimulation (**Figure 6B**) in WT compared to PPARγ-deficient macrophages. However, no differences were reported for *Ankrd29* (**Figure 6D**). As opposed to the dramatic modulation of the kinetics reported under a regulatory microenvironment *in vitro,* the effect observed upon LPS stimulation was limited and gene-specific.

**Figure 6.**
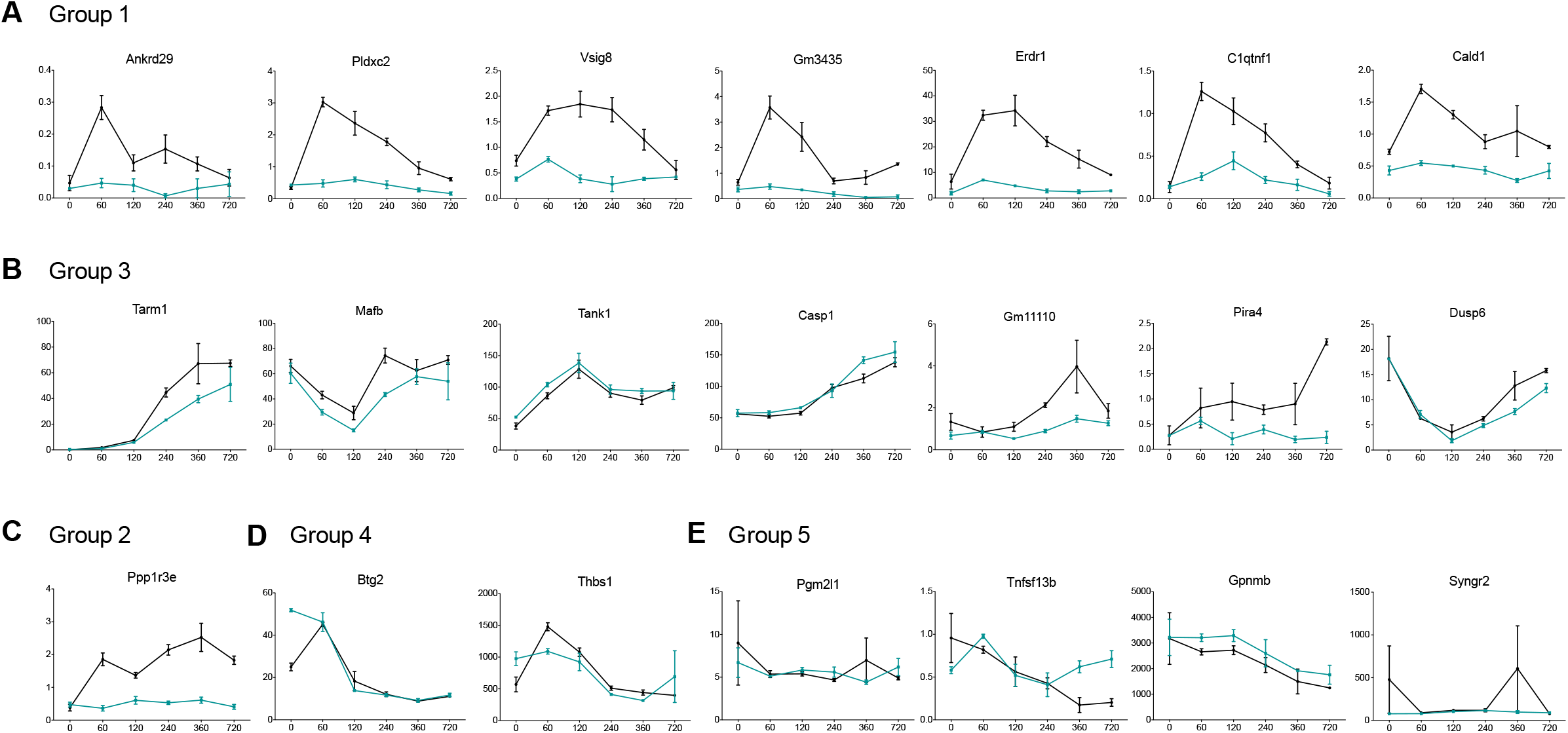
*In vitro* validation of the five selected candidates under pro-inflammatory conditions. WT and LysCre+ BMDM were stimulated with 100 ng/ml LPS and harvested at several time points ranging from 0 to 720 minutes. *Plxdc2* **(A)**, *Ppp1r3e* **(B)**, *Vsig8* **(C)**, *Ankrd29* **(D)**, *C1qtnf1* **(E)**, *TNFα* **(F)**, *Pparγ* **(G)**, and *IL-10* (H) gene expression was measured through qRT-PCR. \**P*-value<0.05 between genotypes, #*P*-value<0.05 within each genotype compared to time 0.

### *Plxdc2* silencing prevents *H.* pylori-induced regulatory phenotype in macrophages and reduces bacterial burden

Once initial screening and validation of the five lead candidates was completed, *Plxdc2* was chosen to further explore its regulatory activity. *Plxdc2* was selected based on the reported expression kinetics under regulatory and pro-inflammatory conditions, together with the fact that *Plxdc2* is a plasma receptor with a known ligand. WT and LysCre+ BMDM were transfected with 20 nM of *Plxdc2* targeted or scrambled siRNA as a negative control using Lipofectamine reagent. BMDM were subjected to the gentamycin protection assay. Cells were harvested at 0, 60, 120, 240 and 360 minutes post-challenge for gene expression analysis, and at 120 minutes for assessment of bacterial loads. Gene silencing resulted in 70% efficiency in WT macrophages, reducing *Plxdc2* expression in this group down to the levels observed in PPARγ-deficient BMDM (**Figure 7A**). At 2 hours post coculture, *Plxdc2* silencing resulted in a 3-fold reduction of *H. pylori* burden in WT macrophages (**Figure 7D**). Additionally, a lower number of *H. pylori* colonies was isolated from both PPARγ-deficient groups in comparison to the WT. However, within LysCre+ macrophages, no differences in bacterial burden were observed after *Plxdc2* suppression via gene silencing. To assess whether the reduced bacterial replication reported in Plxdc2-silenced WT macrophages was due to a potential *Plxdc2* modulation of the macrophage function, we assessed the expression of Arginase 1 (Arg1) and Resistin like alpha (Retnla or fizz1) genes, associated with tissue repair and anti-inflammatory functions in these cells. Both, *Arg1* (**Figure 7B**) and *Retnla* (**Figure 7C**) reported an early response to *H. pylori,* synchronized with the *Plxdc2* expression peak at 60 minutes post-infection in WT Plxdc2-expressing macrophages. However, *Plxdc2* silencing prevented the upregulation of *Arg1* and *Retnla1* after *H. pylori* challenge, leading to a complete abrogation of the 60-minute peak. Similar to the results from bacterial loads, no differences were detected in PPARγ-deficient BMDM irrespective of the state of *Plxdc2* silencing. Indeed, in both LysCre+ groups *Arg1* and *Retnla* exhibited a constant flat expression, also previously observed in all 5 selected candidates (**Figure 5**). Therefore, *Plxdc2* silencing abrogated *H. pylori-* induced regulatory responses in macrophages, resulting in limited bacterial persistence and replication in WT BMDM.

**Figure 7.**
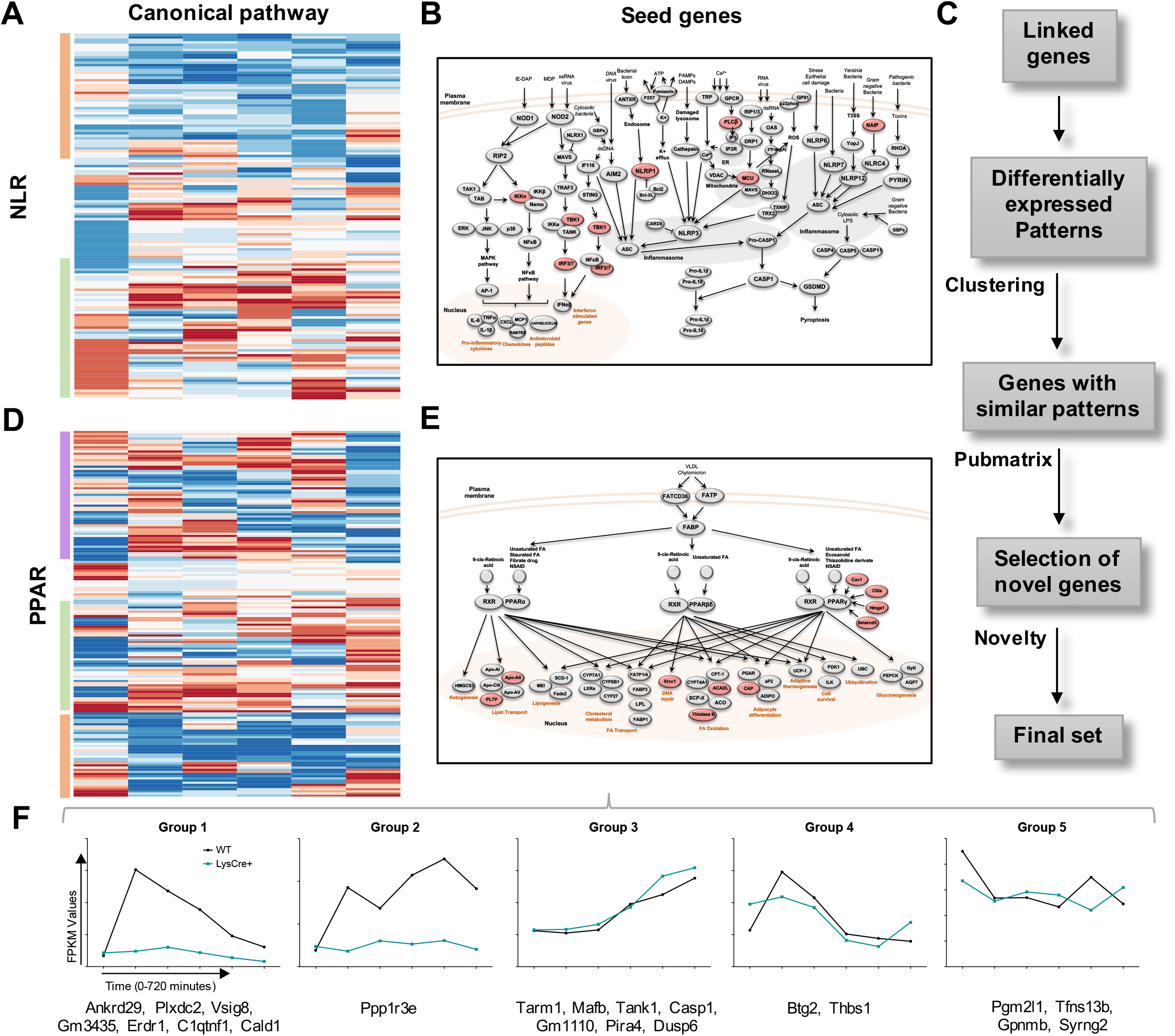
Gene silencing and ligand-induced activation studies to confirm the regulatory role of *Plxdc2.* WT and LysCre+ BMDM were transfected with *Plxdc2* or scrambled siRNA as a negative control prior to *H. pylori* co-culture *in vitro.* Cells were harvested at 0, 60, 120, 240 and 360 minutes post *H. pylori* challenge and *Plxdc2* **(A)**, *Arg1* **(B)**, and *Retnla* **(C)** gene expression was measured. Bacterial burden **(D)** was assessed 120 minutes post co-culture. WT and LysCre+ BMDM transfected with *Plxdc2* or scrambled siRNA as a negative control were treated with the Plxdc2 ligand PEDF at 10nM prior to *H. pylori* challenge. At 120 minutes post coculture cells were harvested, and Arg1 **(E)** and Retnla **(F)** gene expression was assessed. \**P*-value<0.05 between genotypes, #*P*-value<0.05 within treatment.

To further validate the ability of *Plxdc2* to modulate the macrophage phenotype towards a regulatory environment, the *Plxdc2* molecular pathway was activated through the administration of pigment epithelium-derived factor (PEDF), a known Plxdc2 ligand. Interestingly, PEDF treatment slightly increased *Arg1* (**Figure 7E**) and *Retnla* (**Figure 7F**) gene expression in WT and LysCre+ macrophages compared to their respective untreated controls. Indeed, the significant downregulation due to the loss of PPARγ in the untreated group was abrogated after PEDF treatment, exhibiting no differences in comparison to the WT control. Additionally, *Arg1* and *Retnla* expression levels were inhibited in all 4 groups transfected with *Plxdc2* siRNA, reporting no PEDF effects. Therefore, PLXDC2 activation through PEDF treatment increased the expression of anti-inflammatory genes in macrophages and rescued the inflammatory phenotype observed in PPARγ-deficient cells.

## Discussion

We present a novel *ex vivo* co-culture system that leverages the regulatory responses induced by *H. pylori* in macrophages in combination with bioinformatics analyses to discover potential new regulatory genes with immunomodulatory properties based on expression kinetics in WT and PPARγ-deficient BMDM. This new platform was used to identify five genes, *Plxdc2, Ppp1r3e, Vsig8, Ankrd29* and *C1qtnf1* with potential regulatory functions. Induction of regulatory responses through *H. pylori* challenge *in vivo* and *in vitro* confirmed the promising regulatory role of the selected genes. Indeed, the hypothesized regulatory pattern utilized to identify the novel genes, characterized by an early upregulation in WT macrophages and suppression in PPARγ-deficient macrophages, coinciding with the early peak of bacterial burden, was observed in all 5 top lead candidates through *H. pylori* validation *in vitro.* Further, *H. pylori* infection *in vivo* resulted in increased gene expression of all 5 top candidates at day 28 post-infection in WT mice, coinciding with the previously identified greatest accumulation of regulatory macrophages in the gastric mucosa during this bacterial infection (Viladomiu et al., 2017). Therefore, functionally, the correlation between the expression of the 5 candidates and the induction of regulatory mechanisms during *H. pylori* infection, support the immunomodulatory role of the selected genes. Moreover, the validation of the co-culture system provides new insights into the application of bioinformatics screening methods for the discovery of novel molecular targets for treating inflammatory and autoimmune diseases.

*H. pylori* establishes life-long, chronic infections in the gastric mucosa characterized by the induction of mixed immune responses. In addition to the effector mechanisms dominated by Th1 and Th17 cells (Bhuiyan et al., 2014; Carbo et al., 2013; Caruso et al., 2008; D’Elios et al., 1997; Shi et al., 2010; Smythies et al., 2000), *H. pylori* induces strong regulatory responses that suppress mucosal inflammation, contribute to tissue integrity maintenance and prevent effective bacterial clearance (Lundgren, Suri-Payer, Enarsson, Svennerholm, & Lundin, 2003; Raghavan, Fredriksson, Svennerholm, Holmgren, & Suri-Payer, 2003; Raghavan & Quiding-Jarbrink, 2012). Historically, *H.* pylori-associated regulatory responses have been mainly attributed to dendritic cells by inhibition of effector T cells and promotion of regulatory T cells (Kabisch et al., 2016; Kaebisch, Mejias-Luque, Prinz, & Gerhard, 2014; Tanaka et al., 2010). Recently, macrophages have been identified as an integral component in the generation of a regulatory gastric microenvironment during this bacterial infection (Leber et al., 2016; Viladomiu et al., 2017). Applying next-generation sequencing (NGS) and bioinformatics analysis, we demonstrated that in the tightly controlled environment of an *ex vivo* co-culture with BMDM, *H. pylori* challenge drastically influence macrophage gene expression profiles. The significant impact observed in macrophage transcriptomics upon bacterial challenge resulted in the induction of regulatory genes in WT macrophages, that is inhibited by the loss of PPARγ and suppresses the effector mechanisms to promote bacterial persistence. Therefore, the synchronized *ex vivo* co-culture with macrophages, together with the utilization of NGS strategies, is a suitable approach to capture the full spectrum of regulatory responses induced by *H. pylori* and to discover novel regulatory mechanisms with a meaningful impact on the modulation of the immune response.

In addition to the combined NGS and bioinformatics analyses methods, in the current study, gene selection is based on the similarities between the expression kinetics of candidate genes and known host genes with validated regulatory functions. We performed a bioinformatics pattern-based analysis utilizing genes with established regulatory functions to identify novel genes with similar characteristics. In our system, the comparison within macrophages with distinctive immunological steady states (WT versus PPARγ-deficient) was utilized to select the initial core of established genes and to validate the role of the new regulatory gene candidates. To identify the predicted regulatory patterns, we utilized the canonical immune pathways NLR and PPAR. The selection criteria were based on pathways with a dominant regulatory role or including both inflammatory and regulatory members, to allow the full spectrum of expression kinetics and perform a suitable selection of expression patterns associated with immunomodulatory functions. PPARs are nuclear receptors that regulate lipid metabolism and exert potent immunomodulatory functions. Indeed, activation of members in the PPAR pathway result in incremented beta-oxidation of lipid metabolism and suppression of the inflammatory response (Varga, Czimmerer, & Nagy, 2011). The NLR family is a main regulator of the initiation of innate immune responses in macrophages and encompasses more than 20 members (Zhong et al., 2013). As expected, NLR pathway includes an important number of inflammation-driven genes involved in the formation of inflammasomes, such as NLRP1, NLRP3 and NLRC4 (He, Hara, & Nunez, 2016; Masters et al., 2012) or associated with the activation of other inflammatory mechanisms, mainly the NF-κB or MAPK pathways, such as NOD1 and NOD2 (Zhong et al., 2013). Additionally, several genes of the NLR family hold immunoregulatory functions linked to suppression of inflammation, including NLRP10 and NLRX1 (Imamura et al., 2010; Leber et al., 2018; Leber et al., 2017; Xia et al., 2011). Therefore, NLRs constitute a highly heterogeneous and diverse family of pattern recognition receptors (PPR). Other pathways essential for the activation and maintenance of the immune response in macrophages were also considered, including the Toll-like receptors (TLR), the nuclear factor kB (NF-kB) and pathways associated to the production of reactive oxygen species (ROS). However, the strong association of such pathways with the initiation and expansion of inflammatory mechanisms with limited pro-regulatory functions in macrophages (Kawai & Akira, 2006; Liu, Zhang, Joo, & Sun, 2017), limited their potential value in our regulatory gene discovery pipeline. Therefore, to perform an initial screening analysis for regulatory genes, NLR and PPAR pathways were ideal candidates.

The hypothesized immunomodulatory role of the selected regulatory gene candidates was further investigated under the induction of inflammatory mechanisms. LPS stimulation *in vitro* resulted in minor alteration of the expression kinetics in both genotypes. While we observe a trend in *Plxdc2* and *Ppp1r3e* gene expression that supports the hypothesized suppression of immunoregulatory gene levels in pro-inflammatory activated macrophages, PPARγ-deficient cells exhibited slightly greater expression of *Vsig8, Ankrd29* and *C1qtnf1.* LPS stimulation induces strong pro-inflammatory cytokine production, including TNF, IL-6, and IL1β, and the generation of classically-activated macrophages (Arango Duque & Descoteaux, 2014). Therefore, the minor LPS effect on the selected candidates suggests that they might not be associated with the induction or maintenance of pro-inflammatory responses. In conclusion, the observed results in both *in vivo* and *in vitro* experiments under the induction of regulatory and pro-inflammatory mechanisms, suggest that *Plxdc2* and *Ppp1r3e* are the two genes with greater immunomodulatory potential and the most suitable candidates for further validation and study.

*Plxdc2* encodes a 350-amino acid plasma membrane protein known to participate in cell proliferation and differentiation control during the development of the nervous system (Miller et al., 2007; Miller-Delaney, Lieberam, Murphy, & Mitchell, 2011). Our results suggest that *Plxdc2* silencing alters the phenotype of WT macrophages *in vitro* causing a shift towards a more pro-inflammatory state, leading to a significant decrease of bacterial burden and downregulated expression of tissue healing and anti-inflammatory associated genes in comparison to the control group. *Plxdc2,* together with its homologue gene *Plxdc1,* has been identified as one of the membrane receptors of pigment epithelial-derived factor (PEDF) (Cheng et al., 2014), a strong, endogenous anti-angiogenesis factor (Dawson et al., 1999). Interestingly, PEDF treatment modulates macrophage activation in Raw 264.6 cells through the increase of IL-10 production with potential association with PPARγ activation (Yang, Chen, Wu, Ho, & Tsao, 2010; Zamiri, Masli, Streilein, & Taylor, 2006). Cheng et al., reported that PEDF-induced IL-10 production in Raw 264.7 macrophages is dependent on both Plxdc1 and Plxdc2 (Cheng et al., 2014). Our data suggest that *Plxdc2* can induce immunological changes independently of the presence of PEDF in addition to in response to PEDF stimulation. The changes observed in the dynamics of *Plxdc2* over the time course and after silencing of the gene suggest additional yet to be discovered signaling mechanisms with impact on anti-bacterial and overall immune responses. As a cell surface receptor, Plxdc2 is a promising therapeutic target for autoimmune diseases.

The established regulatory effect of PEDF in macrophages, together with the ability to perform ligand-induced activation in addition to the loss of function analysis, triggered the selection of *Plxdc2* for further validation in this initial study. However, the kinetics of *Ppp1r3e* in the validation studies, support the regulatory role of this gene and encourage its further investigation. *Ppp1r3e* gene encodes a glycogen-targeting subunit of the protein serine/threonine phosphatase of type 1 (PP1), a phosphatase protein involved in the regulation of several cell functions, including gene expression, metabolism and cell death (Ceulemans, Stalmans, & Bollen, 2002; Munro, Ceulemans, Bollen, Diplexcito, & Cohen, 2005). Metabolism is an essential component of the immune response (Leber et al., 2017). Activated cells, including macrophages and dendritic cells, undergo metabolic reprogramming, in which glucose metabolism is increased while oxygen consumption is suppressed, to produce energy at fast speed and initiate the mechanisms required for their activation (Kelly & O’Neill, 2015; Rodriguez-Prados et al., 2010). Glycogen-driving subunits, such as *Ppp1r3e,* increase glycogen production promoting glycogen synthase activity (Ceulemans et al., 2002). Moreover, *Ppp1r3e* expression levels are regulated by insulin and are subjected to a significant suppression in diabetic liver (Munro et al., 2005). Interestingly, glycogen also plays a role in the immunometabolic interplay of immune cells (Thwe et al., 2017). Dendritic cells have pools of stored glycogen that is catabolized during the metabolic shift to increase glucose availability and supply the energy needs during these initial steps of the immune response (Thwe et al., 2017). Moreover, deficiencies in glycogen metabolism alter the proper activation of dendritic cells (Thwe et al., 2017). Others have demonstrated that in activated macrophages with overexpression of the glucose transporter GLUT1 and increased glucose consumption, glycogen synthesis is upregulated (Freemerman et al., 2014). Therefore, through the modulation of glycogen metabolism, *Ppp1r3e* might modulate the activation of immune cells, and as a consequence, exert important functions associated with the shape of the immune response.

This study has identified five candidate therapeutic targets with promising host regulatory and immunometabolic roles. Validation studies supported *Plxdc2* and *Ppp1r3e* as two genes with strong regulatory functions and great potential to modulate immune and host responses. Our screening platform can provide new insights in the identification of novel therapeutic targets for the modulation of the immune response that can drive the next wave of the first-in-class therapeutics for widespread and debilitating autoimmune diseases.

## Materials and Methods

### Animal housing and ethic statement

C57Bl/6J wild-type (WT) and PPARγfl/fl;LysCre+ (LysCre+) mice were used for these studies. Mice were gender and age-paired between groups to avoid any external variability. Group allocation was randomly performed. Sample-size estimation was performed using the resource equation approach. LysCre+ were generated by breeding of PPARγ fl/fl into Lys-Cre mice, to produce animals lacking PPARγ in myeloid cells. All mice were bred and housed in the same colony at Virginia Tech in ventilated racks and under a 12-hour light cycle. All experimental procedures performed were approved by the Institutional Animal Care and Use Committee (IACUC) and met or exceeded requirements of the Public Health Service/National Institutes of Health and Animal Welfare Act.

### Bone marrow-derived macrophages (BMDM) isolation and culture

Bone marrow-derived macrophages (BMDM) were isolated as previously described (Zhang, Goncalves, & Mosser, 2008). Briefly, the femur and tibia were excised, cleaned from the attached muscle and sterilized with 70% ethanol. The distal ends of the bones were cut and the bone marrow (BM) flushed out with cold cRPMI_M, containing RPMI 1640 (Corning Incorporated, Corning, NY), 10% Fetal Bovine Serum (Corning Incorporated, Corning, NY), 2.5% Hepes (Corning Incorporated, Corning, NY), 1% Sodium pyruvate (Corning Incorporated, Corning, NY), 1% L-glutamine (Corning Incorporated, Corning, NY), 1% Penicilin/Streptamicin (Corning Incorporated, Corning, NY) and 50uM β-mercaptoethanol (Sigma Aldrich, St. Louis, MO). Osmotic lysis was used to remove red blood cells. Samples were adjusted to 7.5×10^5^ cells/mL with cold cRPMI_M supplemented with 25 ng/mL of recombinant mouse colony-stimulating factor (m-csf, Peprotech, Rocky Hill, NJ) and cultured at 37°C, 5% CO_2_ and 95% humidity to allow their differentiation. At day 3 fresh m-csf-supplemented media was added. On day 6, plates were washed to remove non-adherent cells and BMDM were harvested. Cells were re-suspended in cRPMI_M and seeded in triplicate in 12-well plates (5×10^5^ cells per well). Cells were left to adhere overnight at 37°C, 5% CO_2_ and 95% humidity.

### *H. pylori* culture and preparation of the inoculum

This study was performed using *H. pylori* SS1 strain. *H. pylori* was cultured at 37°C under microaerophilic conditions in Difco Columbia blood agar (BD Biosciences, San Jose, CA) plates supplemented with 7% of Horse laked blood (Lampire Biological Laboratories, Pipersville, PA) and *H. pylori* selective supplement (5mg of Vancomycin, 2.5mg of Trimethoprim, 2.5mg of Cefsulodin and 2.5mg of Amphotericin B, Oxoid, Altrincham, England).

For *in vivo* inoculum preparation, *H. pylori* was harvested at room temperature sterile 1X PBS and adjusted to 2.5×10^8^ colony forming units (cfu) per mL. To obtain the desired concentration, *H. pylori* was adjusted to an optimal density (OD) of 1.2 at a 600-nm wavelength. The association of OD and cfu/mL was based on a previous growth curve that correlated OD with *H. pylori* colony counts. For *in vitro* inoculum preparation, *H. pylori* was harvested in antibiotic-free cRPMI and adjusted to 1×10^8^ cfu/mL as described above.

### Gentamycin protection assay

BMDM cells were washed with 1X PBS and fresh antibiotic-free cRPMI was added to the plates. Cells were infected with *H. pylori* SS1 at a MOI 10 and synchronized by a quick spin to ensure immediate contact. After a 15-minute incubation, non-internalized bacteria were killed by thoroughly washing the cells with PBS/5% FBS containing 100ng/ml Gentamycin (Fisher Scientific, Pittsburg, PA). Cells for time-point 0 were washed with PBS and immediately collected for downstream assays. The cells allocated for the remaining time points (30 min to 12 hours) were covered with culture media until collection for bacterial re-isolation or assessment of differential gene expression by qRT-PCR and whole transcriptome analyses. Each validation study included between two and five biological replicates and was repeated at least twice.

### Bacterial Re-isolation from BMDM

BMDM cells were washed 3 times with sterile 1X PBS. 200uL of Brucella broth (BD Biosciences, San Jose, CA) were added to the well and cells were detached using a cell scraper. Cell suspensions were sonicated 5 seconds to release intracellular bacteria. Serial 10-fold dilutions of the original homogenate were plated into the *H. pylori* plates described above. After for 4 days of culture at 37°C under microaerophilic conditions, colonies were counted.

### Gene Expression

BMDM cells were collected in ice-cold RLT (supplemented with β-mercaptoethanol) and stored at – 80°C until RNA isolation was performed. Following mouse euthanasia, stomachs were excised from the animal and longitudinally opened. Collected tissues were rinsed twice in 1X PBS and stored in 350uL of RNAlater (Fisher Scientific, Pittsburg, PA) at −80°C. Total RNA was extracted from BMDM and stomach using the RNeasy mini kit (Qiagen, Hilden, Germany) following manufacturer’s instructions. RNA concentrations were quantified with a nandrop (Invitrogen, Carlsbad, CA) at 260nm. iScript cDNA synthesis kit (Bio-Rad, Hercules, CA) was utilized to generate cDNA from RNA samples. Primer-specific amplicons were produced by PCR using Taq Polymerase (Promega, Madison, WI), followed by a purification step using the Mini-Elute PCR purification kit (Qiagen, Hilden, Germany). Standard curves were generated by a series of dilutions from a known concentration of the purified primer-specific amplicon, starting at 1×10^6^pg. Total gene expression levels were assessed through a quantitative real-time PCR (qRT-PCR) using a CFX96 Thermal Cycler (Bio-Rad, Hercules, CA) and SYBR Green Supermix (Bio-Rad, Hercules, CA). βeta-actin expression was utilized to normalize the expression levels of targeted genes. Primer sequences are included in supplementary information.

### Global Transcriptome analysis

RNA isolated from WT and LysCre+ BMDM collected at time-points 0, 30, 60, 120, 240, 360 and 720 minutes post-infection was submitted for whole transcriptome gene expression analysis using Illumina Hiseq (Biocomplexity Institute of Virginia Tech Core Lab Facilities). The biological replicates for each sample were three. A replicate for three of the samples (LysCre+ HP 0 minutes, LysCre+ HP 120 minutes and WT uninfected 240 minutes) did not pass quality control, therefore these three samples consisted of 2 biological replicates. Once fastq files containing 100bp-long pair-end reads were received, poor quality reads (>40% of bases with PHRED score <10; percentage of N greater than 5%; and polyA reads) were excluded. Through the utilization of Bowtie (Langmead, Trapnell, Pop, & Salzberg, 2009) (version: 1.0.0) with parameters set to −l 25 −I 1 −X 1000 −a −m 200’, the remaining reads we mapped to RefSeq (mm10 from http://genome.ucsc.edu/). To calculate gene expression levels we used RSEM (Li & Dewey, 2011), a program based on expectation-maximization algorithm. FPKM (Trapnell et al., 2010) (fragments per kilobase per million sequenced reads) was used as the measurement of expression level. Data were submitted to NCBI’s GEO database (Accession Number GSE67270).

### Bioinformatics analysis

As described in **Figure 3C**, to analyze the RNAseq data, an initial dataset of genes linked to the selected NLR and PPAR candidates was generated. Briefly, the Genome-scale Integrated Analysis of gene Networks in Tissues (GIANT) and Gene Expression Omnibus (GEO) databases were used and integrated in the Ingenuity Pathway Analysis (IPA) software to build the initial group of genes. Hierarchical based clustering was employed to obtain differentially expressed patterns within the combined initial NLR and PPAR genes and the linked dataset generated. The *hclust* method with Ward’s minimum variance method and Manhattan distance metric in R were used to cluster the data. Another clustering cycle was performed in order to obtain a larger set of genes with similar patterns of interest. The generated dataset was enhanced for novelty through an abstract searching using the Pubmatrix tool, and a final dataset of genes was obtained.

### Gene silencing and BMDM treatments

For gene silencing studies, WT and LysCre+ BMDM were transfected with 20nM siRNA (27mer Dicer-substrate siRNA, DsiRNA, for the targeted gene or scrambled sequence as a negative control, Integrated Device Technology, San Jose, CA) using Lipofectamine RNAiMax Transfection Reagent (Thermo Fisher Scientific, Waltham, WA) 48 hours before the infection. Media was replaced 6 hours post-transfection. Cells were exposed to *H. pylori* following the gentamycin protection assay described above. Gene knock-down was validated by qRT-PCR.

For the induction of a pro-inflammatory environment during gentamycin protection assay, BMDM were pre-treated with LPS (100ng/mL, Sigma Aldrich, St. Louis, MO) and rIFNγ (100ng/mL) overnight. For the LPS validation, BMDM were treated with LPS (100ng/mL) for 1, 2, 4, 6 and 12 hours. For Plxdc2 activation, 48 hours post siRNA transfection, WT and LysCre+ BMDM were treated with PEDF (10nM) for 24hours. Then, BMDM were subjected to gentamycin protection assay.

### *In vivo H. pylori* infection

8 to 12-week-old WT and LysCre+ mice were transferred to an ABSL2 room in the same colony at Virginia Tech in a ventilated rack and under a 12-hour light cycle. On days 0 and 2 of the project, mice were administered 200uL of freshly prepared 5×10^7^ cfu of *H. pylori* SS1 in 1X PBS through orogastric gavage. These studies also included a non-infected group administered with 200uL of 1X PBS without bacteria. Mice were monitored for signs of disease weekly and stomach samples were collected at 28 days post-infection.

### Statistics

Data are expressed as mean values and standard error of the mean represented in error bars. To calculate the significance of the RNAseq dataset from the global transcriptome analysis, all genes with median expression level in all samples greater than 0 were included in a 3-way (genotype, treatment and time) ANOVA analysis. Normal quantile transformation (qqnorm from R (Ihaka & Gentleman, 1996)) was used to normalize the FPKM to fit the normality assumption of ANOVA (tested with Kolmogorov-Smirnov test). The 3-way ANOVA analysis was carried in R (Ihaka & Gentleman, 1996), FDR (Benjamini & Hochberg, 1995) and Bonferroni were used to calculate the adjusted P-values. To determine significance of the standard data, Analysis of variance (ANOVA) was performed using the general linear model procedure in SAS (SAS Institute). Significance was considered at p ≤ 0.05 and significant differences were identified with an asterisk (genotype) or pound sign (infection or treatment).

## Legends of Figure Supplements

**Figure 2 – Figure supplement 1.**
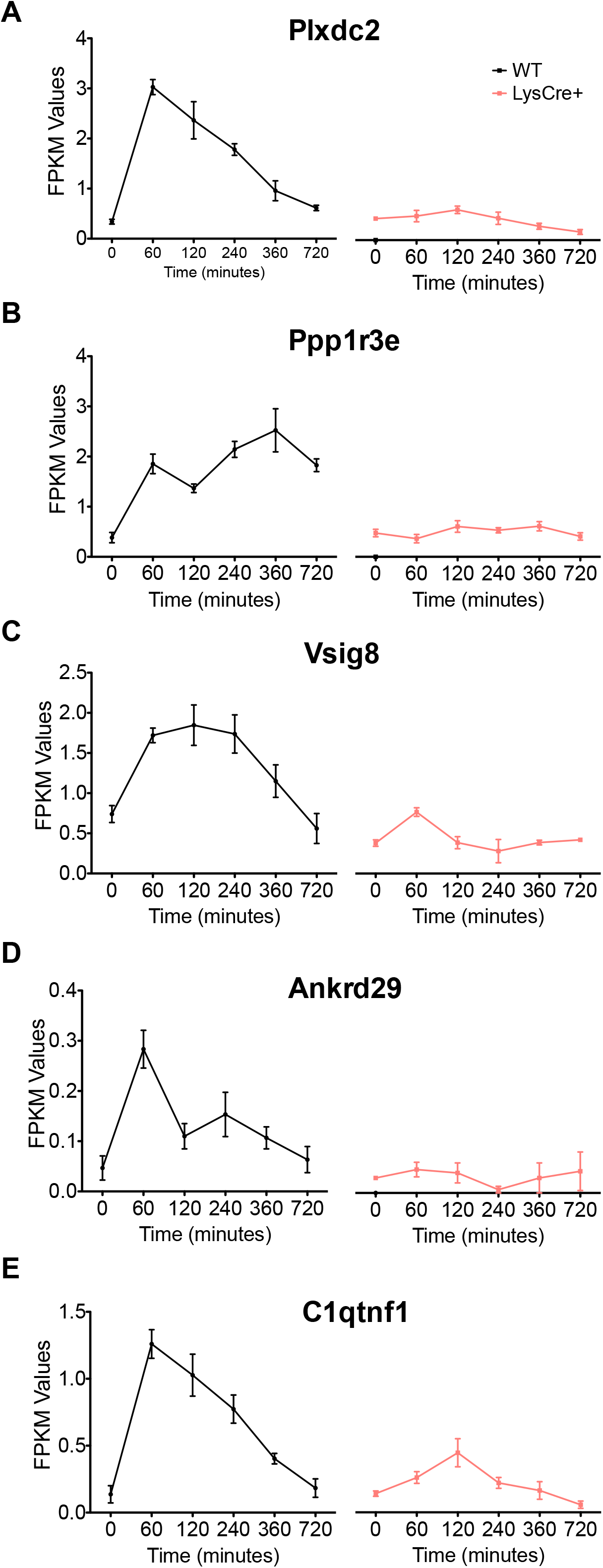
Expression kinetics of several pro-inflammatory genes during *Helicobacter pylori* co-culture. Plots represent the RNAseq reads comparing WT and LysCre+ BMDM in the distinctive time points of the gentamycin protection assay of *MCP1* **(A)**, *MCP5* **(B)**, *IL6* **(C)**, *IFNγ* **(D)**, *IL12a* **(E)**, *IL12b* **(F)**, *CXCL1* **(G)**, *CXCL10* **(H)**, and *MIP-1α* **(I)**.

**Figure 2 – Figure supplement 2.**
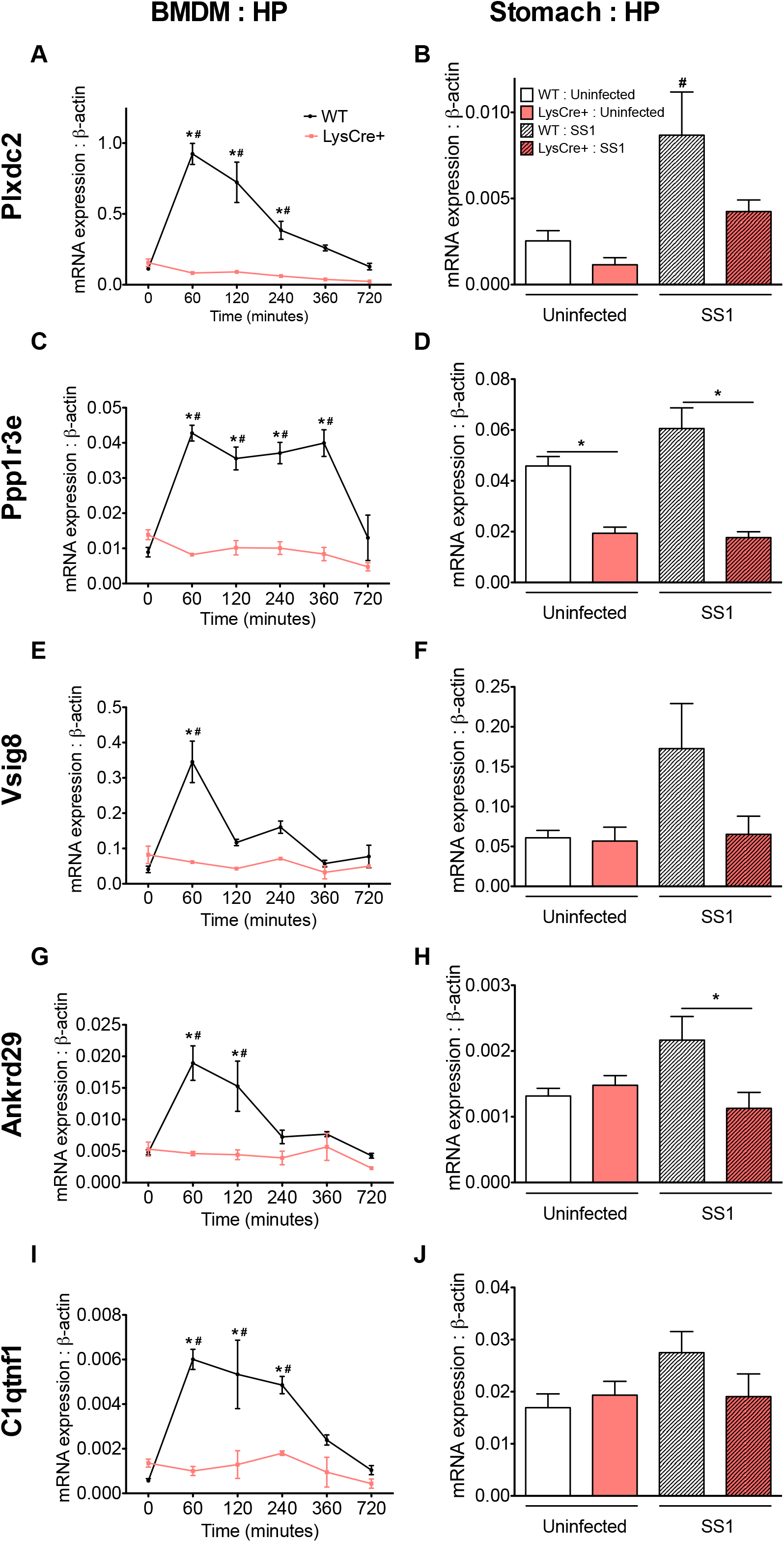
3-way ANOVA analyses revealed eight genes with significant differential expression pattern. Plots represent the RNAseq values of *Chil1* **(A)**, *Etv5* **(B)**, *ligpi* **(C)**, *Ptger4* **(D)**, *Sqle* **(E)**, *Osm* **(F)**, *Rptoros* **(G)**, and *Hspa2* **(H)**. *P*-value<0.05.

**Figure 2 – Figure supplement 3.**
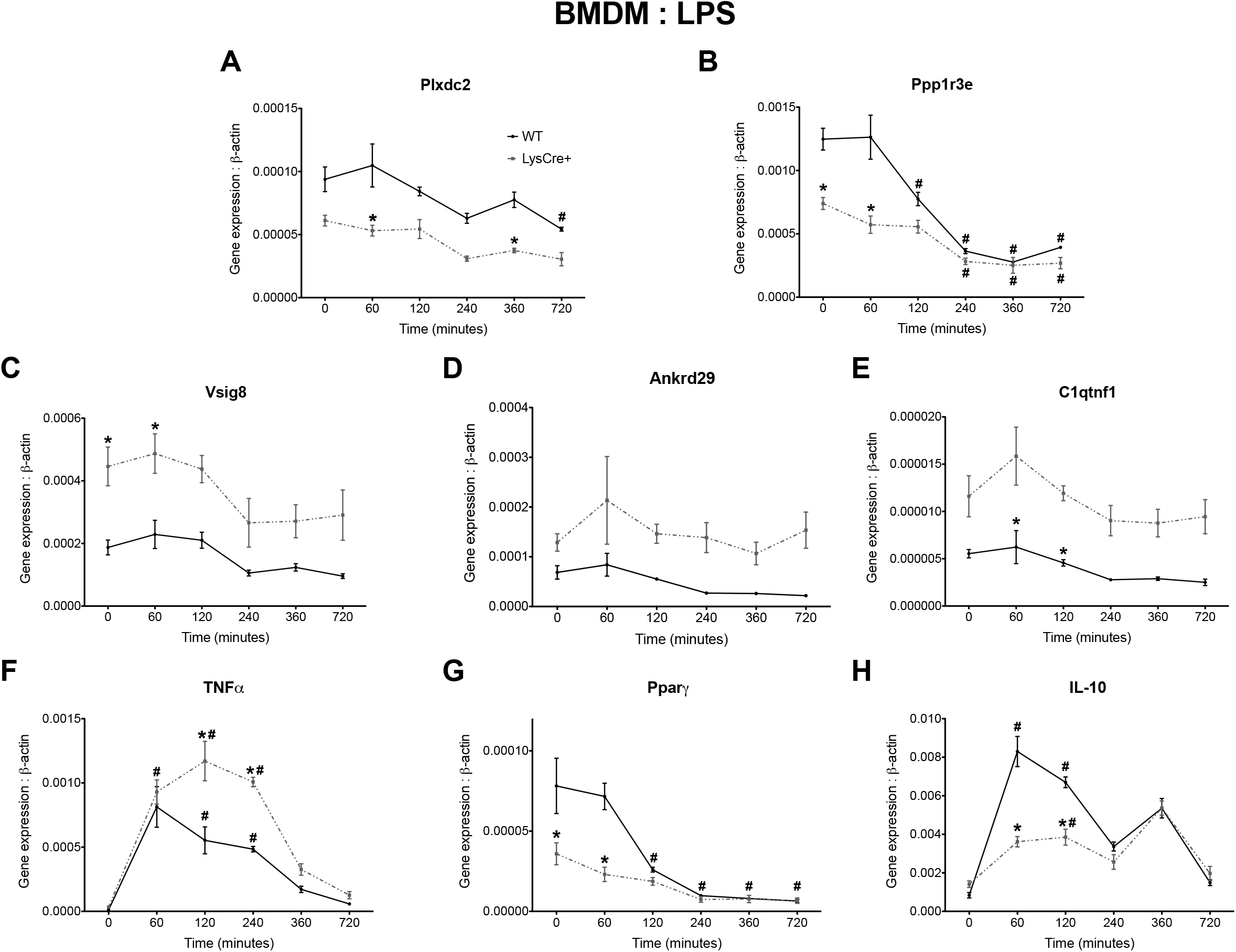
Validation of *Chill* and *ligp1* gene silencing by qRT-PCR. WT and LysCre+ macrophages were transfected with *Chil1*-targeted, *ligpi*-targeted or negative scrambled siRNA prior to *H. pylori* challenge. Cells were harvested 120 minutes after *H. pylori* co-culture. *Chil1* **(A)** and *ligpi* **(B)** gene expression was assessed. #*P*-value<0.05 within treatments.

**Figure 3 – Figure supplement 1.**
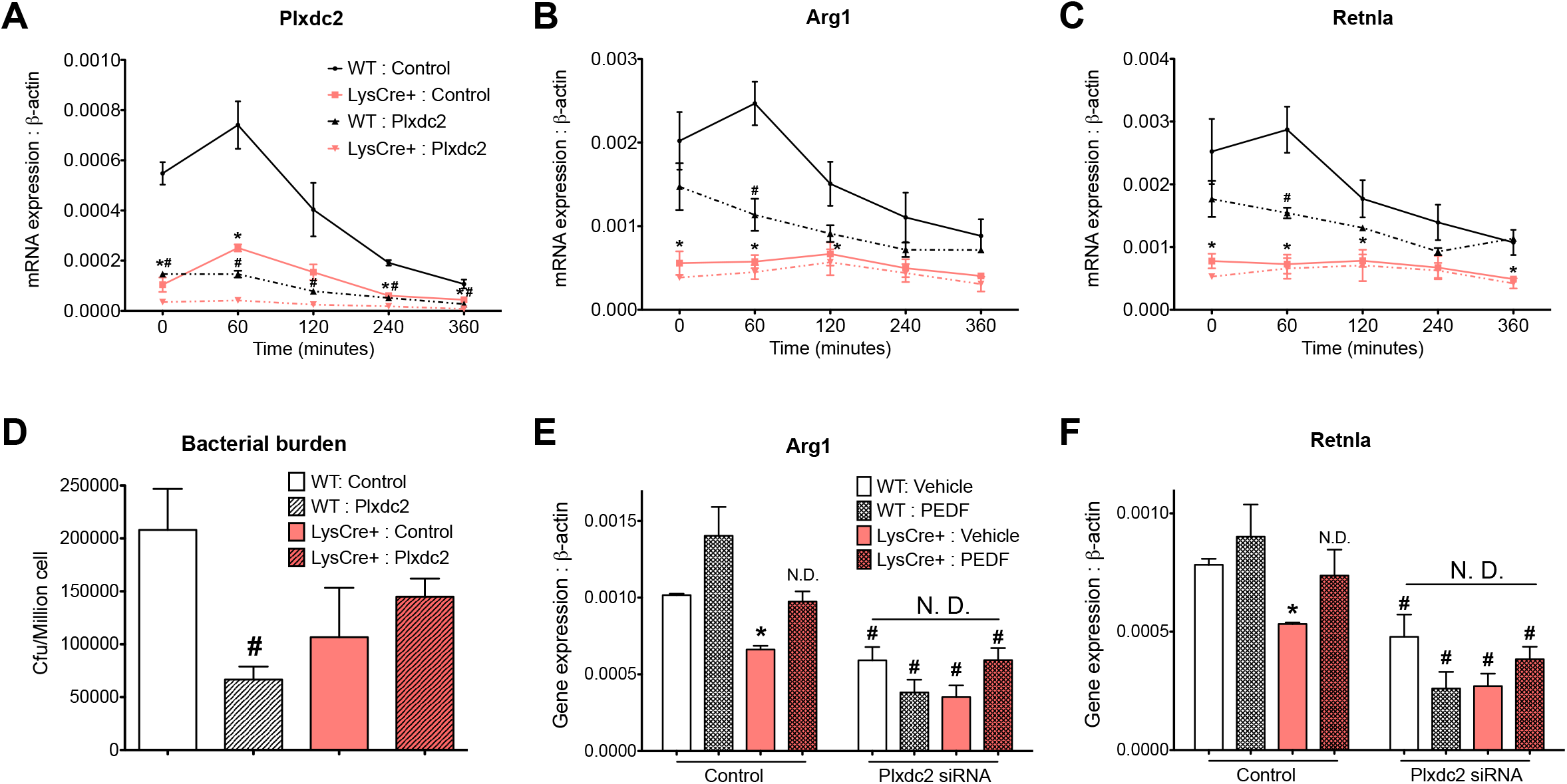
Final set of genes from the bioinformatics analysis consists of 21 candidates classified in five groups based on the expression kinetics. Plots represent the RNAseq reads at each time point of the experiment comparing WT and LysCre+ BMDM. Group 1 **(A)** includes *Ankrd29, Plxdc2, Vsig8, Gm3435, Erdr1, C1qtnf1,* and *Cald1.* Group 3 **(B)** consists of *Term1, Mafb, Tank1, Casp1, Gm11110, Pira4,* and *Dusp6.* Group 2 (C) only includes *Ppp1r3e,* whereas group 4 **(D)** includes *Btg2* and *Thbs1.* Group 5 **(E)** consists of *Pgm2l1, Tnfsf13b, Gpnmb,* and *Syngr2.*

## Supplementary Files

Supplementary File 1. qRT-PCR primers utilized in this study.

